# Electrophysiological correlates of proactive and reactive inhibition in a modified visual Go/NoGo task

**DOI:** 10.1101/2021.07.12.451610

**Authors:** Jiří Kutý, Alena Damborská, Pavla Linhartová, Martin Lamoš, Barbora Jeřábková, Daniela Rudišinová, Martin Bareš, Tomáš Kašpárek

**Affiliations:** Department of Psychiatry, Faculty of Medicine, Masaryk University, Brno, Czech Republic; CEITEC - Central European Institute of Technology, Masaryk University, Brno, Czech Republic; 1st Department of Neurology, St. Anne’s Faculty Hospital, Masaryk University, Brno, Czech Republic; Department of Neurology, School of Medicine, University of Minnesota, Minneapolis, MN, USA

**Keywords:** Borderline personality disorder, impulsivity, Go/NoGo task, EEG, proactive inhibition, reactive inhibition

## Abstract

Impulse control is crucial for everyday functioning in modern society. People with borderline personality disorder (BPD) suffer from impulse control impairment. According to the theory of dual mechanisms of control, motor impulse control can be divided into proactive and reactive modes. Proactive inhibition is involved before an event that might require inhibitory control. Reactive inhibition is initiated after the occurrence of an event that requires inhibitory control. Few studies have focused on proactive inhibition in relation to impaired impulse control, moreover electrophysiological evidence is scarce. Therefore, in the search for electrophysiological correlates of proactive and reactive inhibitions, we assessed event-related potentials elicited during a modified emotionally neutral visual Go/NoGo task in 28 clinically impulsive BPD patients and 35 healthy control (HC) subjects. In both groups, proactive inhibition was associated with enhanced late prestimulus activity and a suppressed poststimulus N2 component. In both groups, reactive inhibition was associated with enhanced poststimulus N2 and P3 components. We found no electrophysiological differences between HC subjects and BPD patients and both groups performed similarly in the task. Hence, the clinically observed impulse control impairment in BPD might act through different mechanisms other than altered inhibitory control in an emotionally neutral task.

## 1. Introduction

Every day we face a multitude of decisions, which impact our lives to a lesser or greater degree. Should I choose a tasty but rather unhealthy meal, or instead opt for a healthier but less appetizing alternative? Do I really need to buy a new jacket, even if I have one like it already in my closet? Choosing the more immediately gratifying alternative, acting on the spur of the moment, is considered a sign of impulsive behavior. While most healthy individuals sometimes engage in impulsive behavior, extensive impulsivity is deleterious not only to one’s subjective well-being but also to the well-being of broader society. Clinically, impulsivity refers to the inability of an individual to inhibit undesired thoughts or behaviors. Impulsivity can be manifested in a broad range of pathological behaviors like substance abuse, gambling, dangerous sexual behavior, risky driving, binge eating, aggression, and self-harm (Barteček et al., 2019; Reeves-Dudley, 2017; Cyders et al., 2009). Impulsive traits are characteristic of several neuropsychiatric diseases. For borderline personality disorder (BPD), impulsivity is a core defining feature.

With a lifetime prevalence of 1.4 – 5.9% (Lenzenweger et al., 2007; Grant et al., 2008) and high levels of suicidal behavior (Pompili et al., 2005; Black et al., 2004), BPD is a severe and potentially life-threatening disorder. There is a long-established link between impulsivity levels and the occurrence of suicidal behavior in BPD (Brodsky et al., 1997; Links et al., 1999; Soloff, 2000; Rihmer et al., 2010). Inhibitory mechanisms, responsible for preventing impulsive actions are thought to be impaired in BPD patients, leading to dangerous, unpredictable behavior.

According to the dual mechanisms of cognitive control theory (Braver, 2012), inhibitory mechanisms can be divided into proactive and reactive modes. Proactive inhibition is involved prior to an event that might require inhibitory control. It is initiated for example by a warning sign informing a person to watch out for cars while crossing the road. It prepares for the motor inhibition that might be required later. Reactive inhibition, on the other hand, is an active inhibitory reaction driven by an external stimulus. It is present when a person abruptly stops during a road crossing after having been startled by the loud noise of a car horn. While proactive inhibition is considered to be a top-down regulatory endogenous process, reactive inhibition is viewed as a bottom-up exogenous process (Liebrand et al., 2017; Zandbelt, 2011).

Several cognitive-motor tasks have been employed to study inhibitory mechanisms (Linhartová et al., 2021), most notably the Go/NoGo task. In this task, inhibition is usually assessed using behavioral indicators like accuracy or reaction time. Additionally, brain imaging techniques such as electroencephalography can be used to study electrophysiological correlates of task-related brain activity. Particularly, event-related potentials (ERPs) have been widely utilized to study mental activity during the Go/NoGo task. Two major ERP components traditionally associated with inhibition are the N2 and P3 (Kok et al., 2004).

The N2, a negative-going wave emerging 200 – 300 ms after the stimulus onset, was suggested to reflect the top-down mechanisms responsible for inhibiting the tendency to respond (Kaiser et al., 2003). Nevertheless, this component was previously associated with conflict monitoring (Donkers and van Boxtel, 2004) and more recently interpreted as an index for early cognitive processes necessary in implementing inhibitory control rather than the actual inhibitory brake (Luijten et al., 2014).

The P3 component, a positive-going wave emerging 260 – 450 ms after the stimulus onset (Sutton et al., 1965; Galdo-Alvarez et al., 2016), is arguably the strongest electrophysiological indicator of inhibition. A higher P3 amplitude (Nguyen et al., 2016; Wessel and Aron, 2015) and a shorter peak latency (Roche et al., 2005) were observed in successful compared to unsuccessful inhibitions during inhibition tasks, suggesting slower unfolding of inhibitory processes in unsuccessful inhibition. The P3 thus appears to reflect the processes closely related to inhibition of the motor system (Kenemans, 2015; Kok et al., 2004; Dimoska et al., 2006; Band and van Boxtel; 1999). Impaired inhibition in BPD patients was previously shown to be reflected in a reduced NoGo-P3 amplitude (Ruchsow et al., 2008). Albert et al. (2019) found lower frontocentral P3 in BPD subjects in both Go and NoGo trials during the Go/NoGo task.

The classical Go/NoGo task does not allow, however, the distinction between proactive and reactive inhibitory processes. Recently, modified versions of the Go/NoGo task have been proposed to reveal ERP correlates of both proactive and reactive inhibitory processes in a healthy population. For example, Liebrand et al. (2017) used a cued version of the Go/NoGo task. In that study the authors associated proactive inhibition with an increased contingent negative variation (CNV) component in the cue period and an increased N1 component in the poststimulus period. In order to study proactive inhibitory processes in healthy subjects, Albares et al. (2014), extended the classical version of the Go/Nogo task with a “certain” Go condition. The authors used a blind source separation technique and found a specific independent component originating in the supplementary motor area approximately 170ms after the stimulus onset. Its amplitude differed in conditions with and without uncertainty, i.e. conditions requiring and not requiring proactive inhibition. The authors associated this independent component with both automatic response activation and concurrent automatic inhibition.

Only a few studies have focused on proactive inhibition in relation to impaired impulse control (Sharif-Razi et al., 2019; Van Hulst et al., 2018; Castro-Meneses et al., 2015), and none of them have investigated this relationship using an electrophysiological approach. Therefore, in the current study we aimed to assess ERP brain activity in clinically impulsive BPD patients using the modified Go/NoGo task (Albares et al. 2014). In the uncertain variant of this task, two different responses are required - either to press a button or to refrain from any motor action. In the certain task variant, motor action is required in response to every stimulus. Thus, we expected proactive inhibition to be involved in the uncertain but not in the certain task variant. Similarly, within the uncertain task variant, we expected reactive inhibition to occur only in the NoGo condition where no motor action was required.

Contrary to the original study (Albares et al. 2014), we used global electrophysiological measures of ERP activity such as global field power and global map dissimilarity (Habermann et al., 2018), that allowed for direct comparison with the results of other ERP studies. In the current study, we searched for: (1) global electrophysiological correlates of proactive and reactive inhibitions; (2) the difference in behavioral and electrophysiological measures of proactive and reactive inhibitions between clinically impulsive BPD patients and healthy controls.

## 2. Methods

### 2.1 Subjects

Data from 35 patients diagnosed with borderline personality disorder (BPD) and 40 healthy controls (HC) were collected. BPD patients were recruited from the Department of Psychiatry of the University Hospital Brno, Czech Republic, and through outpatient psychiatrists. HC patients were recruited via an internet advertisement and screened using the Mini international neuropsychiatric interview (Sheehan et al., 1998). Twelve subjects were excluded due to EEG artifacts, resulting in 28 BPD and 35 HC subjects used for the ERP analyses. Both groups were comparable in terms of age in years (BPD mean = 23.48 ± 4.85; HC mean = 23.88 ± 7.06, t = -0.259, p = 0.796) and gender (4 males in both the BPD and HC groups). All subjects in both groups were right-handed. Self-reported impulsiveness was assessed using the UPPS-P scale. The BPD group showed higher self-reported impulsivity on the UPPS-P scale (BPD mean ± SD = 159.72 ± 25.56, HC mean ± SD = 126.62 ± 22.99), t (48.54) = 5.13, p <0.001).

### 2.2 Experimental Protocol

The participants were seated in a dimly illuminated room, facing a computer screen while performing the modified Go/NoGo task. The white capital letters A and B were displayed in the center of a black screen as experimental stimuli. Each stimulus was preceded by a fixation cross that was displayed with a random duration of 2 to 6 s (Figure 1). The stimuli were displayed for 0.2 s, followed by a black screen that appeared for 2 seconds. Thus, the inter-stimulus interval ranged from 4.2 s to 8.2 s. The fixation cross was either red, signaling the uncertain task variant, or green, signaling the certain task variant. In the uncertain task variant, either the letter A (Go stimulus) or B (NoGo stimulus) appeared with an equal probability. In the certain task variant, the letter A (Go stimulus) was always displayed. The red cross was twice as common as the green cross. This resulted in an equal number of the three experimental conditions: Go certain (the letter A following the green cross), Go uncertain (the letter A following the red cross), and NoGo (the letter B following the red cross) conditions.

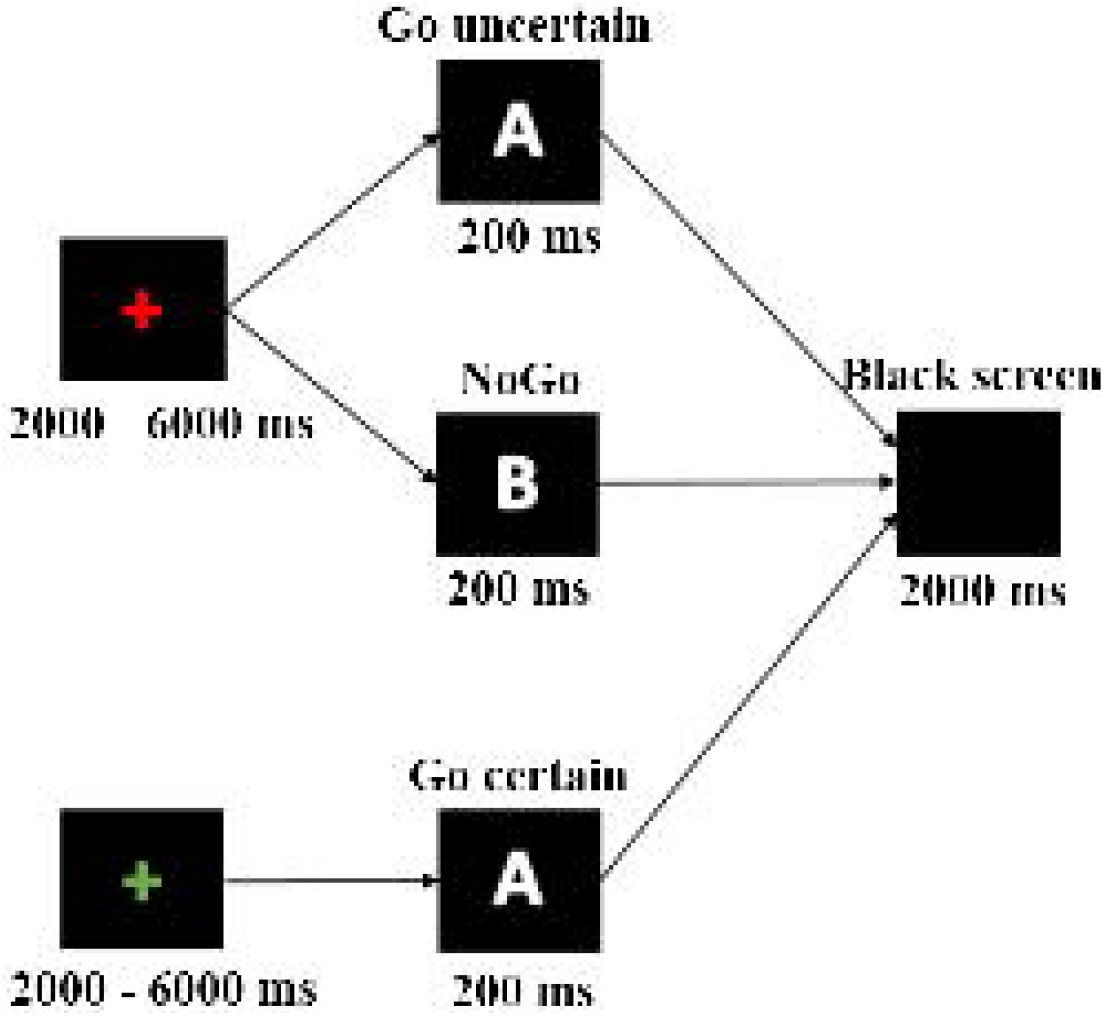

Before the experiment, all participants completed a short practice block to get acquainted with the task. The participants were instructed to press a button as quickly as possible on both the Go certain and Go uncertain stimuli, and to suppress this action when the NoGo stimulus appeared. All participants were presented with 72 stimuli of each type. The task was presented in three identical blocks, with breaks between the blocks being used for adjustment of electrode impedances.

### 2.3 Data acquisition and pre-processing

We used high-density electroencephalography (EEG) to evaluate the temporal and spatial distribution of ERPs as possible electrophysiological correlates of proactive and reactive inhibition. Data were recorded using the EGI 129 electrodes system (EGI GES 200; Electrical Geodesic Inc., OR, USA) referenced relative to a Cz electrode. The signal was digitized at a sampling rate of 1000 Hz; impedances were kept below 50 kΩ. Offline, the data were band-pass filtered from 1 to 40 Hz. Independent component analysis as implemented in the EEGlab (Delorme and Makeig, 2004) was applied to suppress the ballistocardiogram and oculomotor components. The EEG was then downsampled to 500 Hz and the previously identified biased channels were interpolated. The data were then re-referenced to the average reference.

The remaining EEG artifacts were identified by visual inspection using SignalPlant (Plesinger et al., 2016) and discarded after epoching. Pressing the button in response to the NoGo stimulus or failing to press it in response to the Go stimulus were considered commission or omission errors, respectively (Table 1). Only the error-free and artifact-free trials were used for averaging. In each subject, the post-stimulus epochs from 0 to 2000 ms were averaged separately for each condition using the stimulus onset as a trigger. Similarly, the post-fixation-cross epochs from 0 to 2000ms were averaged separately for each task variant.

### 2.4 ERP analysis

Two global tests across all electrodes were used, one to test for global strength difference of the electric field and the other to test for differences in the topography of the potential distribution (Michel and Murray, 2012). This approach to EEG data analysis was successfully used in previous studies (Bailey et al. 2014; Berchio et al., 2017). The difference in map strength was assessed by calculating the Global Field Power (GFP), first introduced by Lehmann and Skrandies (1980). GFP reflects the degree of neuronal synchronization (Skrandies, 1990). It includes the differences between all possible electrode pairs and is equivalent to the topographical standard deviation of the potentials. The topographic ANOVA (TANOVA) was used to find differences in potential scalp distribution between groups and conditions. A TANOVA calculates the topographic difference between two scalp maps (global map dissimilarity). The data for TANOVA analysis were normalized prior to analysis. The difference in normalized scalp maps is considered a qualitative effect whereas the difference in GFP is considered a quantitative effect (Habermann et al., 2018). GFP shows the magnitude of difference regardless of their source in the brain, whereas a TANOVA on normalized data shows whether the spatial distribution, location, and orientation differs between conditions. The epochs 0-2000ms were analyzed using 5000 permutations and an alpha level of 0.001 using Ragu software (Koenig et al., 2011). Additionally, only time-windows longer than 10ms with a p-value below the alpha level were considered significant. We used the permutation analysis for the 2×2 design and posthoc tests to reveal the post-stimulus difference in evoked activity between the Go certain and Go uncertain and between the Go uncertain and NoGo conditions and the two groups.

Similarly, the post-fixation cross differences in evoked activity were analyzed using a 2×2 design and permutation posthoc test. The difference ERP waveforms were computed on an individual level by averaging across the trials followed by subtracting between the different conditions or different task variants. Subsequently, grand averages were constructed from the computed individual waveforms.

### 2.5 Behavioral data analysis

We tested for the difference in reaction times and the number of commission and omission errors using R statistical software (Team RC, 2020) and Jamovi (Şahin and Aybek, 2019). A two-way ANOVA was conducted to test for differences in reaction times. Non-parametric tests were conducted to compare error rates between groups and conditions. Additional tests were conducted between conditions for a statistically significant factor.

## 3. Results

### 3.1 Behavioral results

The task performance was good in both groups. The mean accuracy across subjects exceeded 80% and the mean reaction time was less than 400ms in all conditions both in the BPD and HC groups, (see Table 2). We observed lower accuracy in the BPD than the HC group in the Go uncertain condition, but not in the Go certain or NoGo conditions. In addition, both groups were significantly more accurate in the Go uncertain condition compared to the other two conditions (Fig. 2). The mean reaction time did not significantly differ between groups either in the Go uncertain condition or in the Go certain condition (Table 2). Longer reaction times were observed in the Go uncertain compared to the Go certain conditions in both groups (p < 0.001) (Fig. 3).

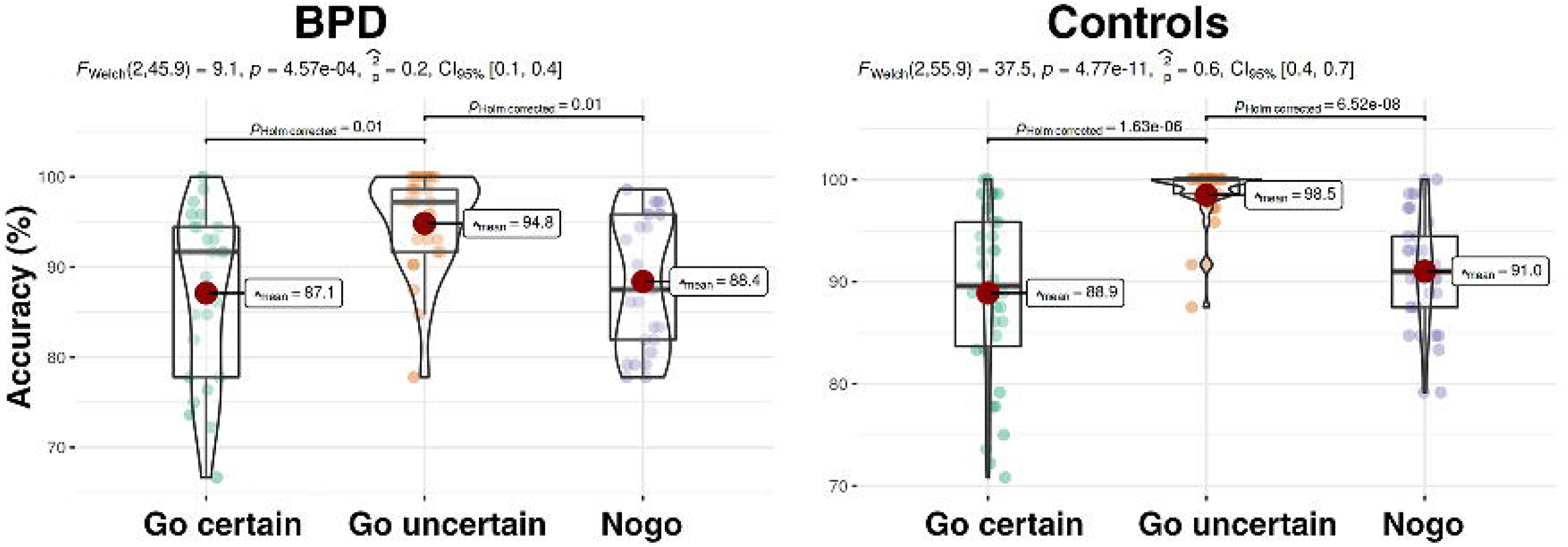

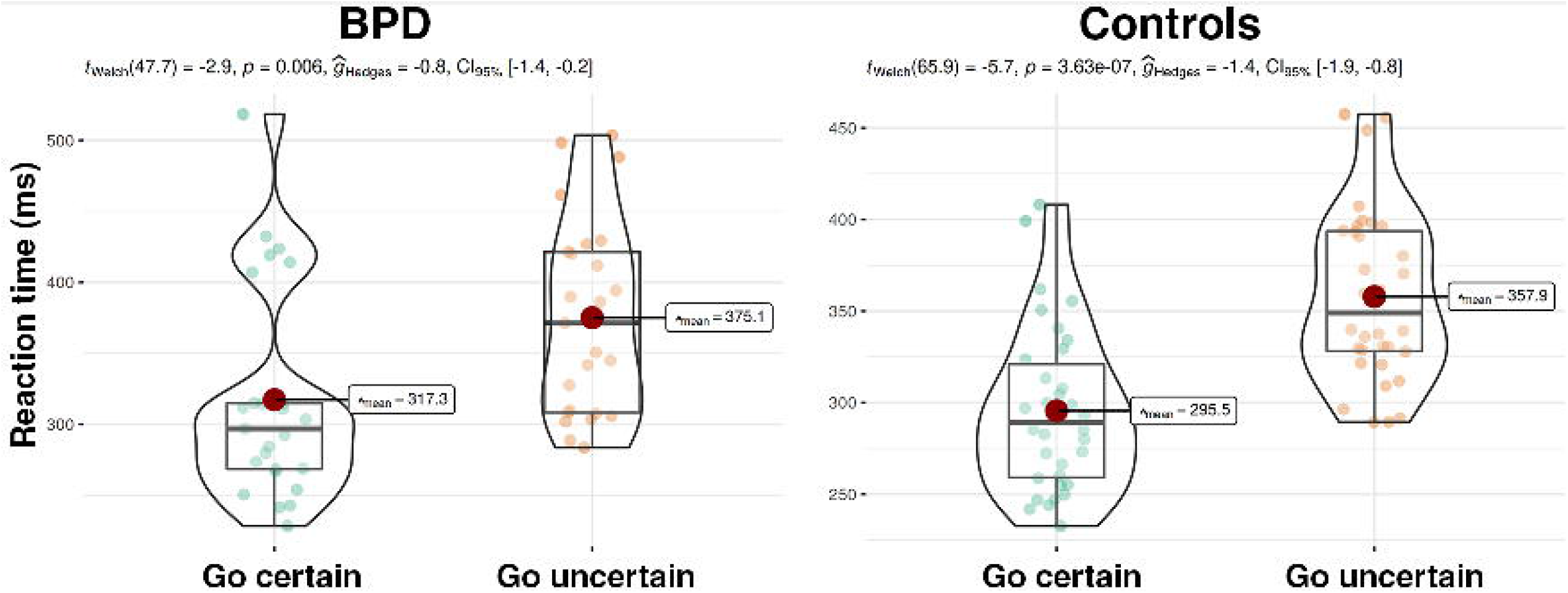

### 3.2 Evoked response results

All conditions both in the patient and control groups elicited stimulus-locked P2, N2, and P3 components within the reaction-time period (Figure 4); grand average waveforms in all electrodes are presented in Figure 5. Moreover, in both groups, we observed a fixation cross-locked activity (Figure 6). The permutation analysis revealed a significant main effect of condition in both GFP and TANOVA metrics. No main effect of group was detected in any metric. Figures 7 and 8 show different waveforms for the conditions in each group. Figure 9 shows different waveforms for the task variants in each group.

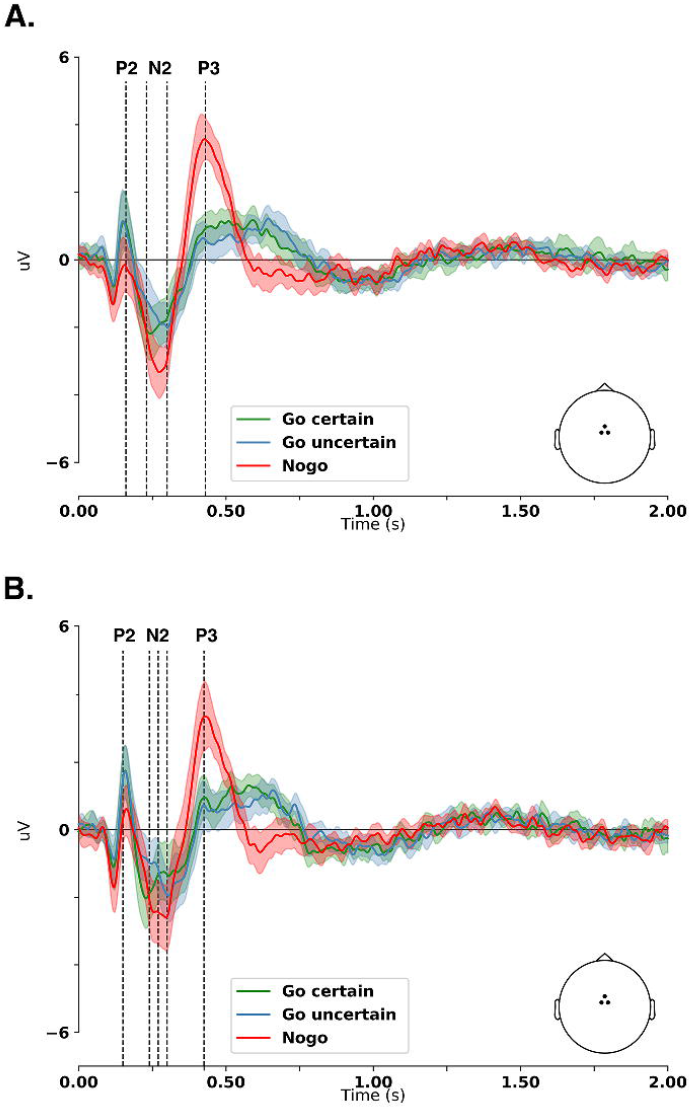

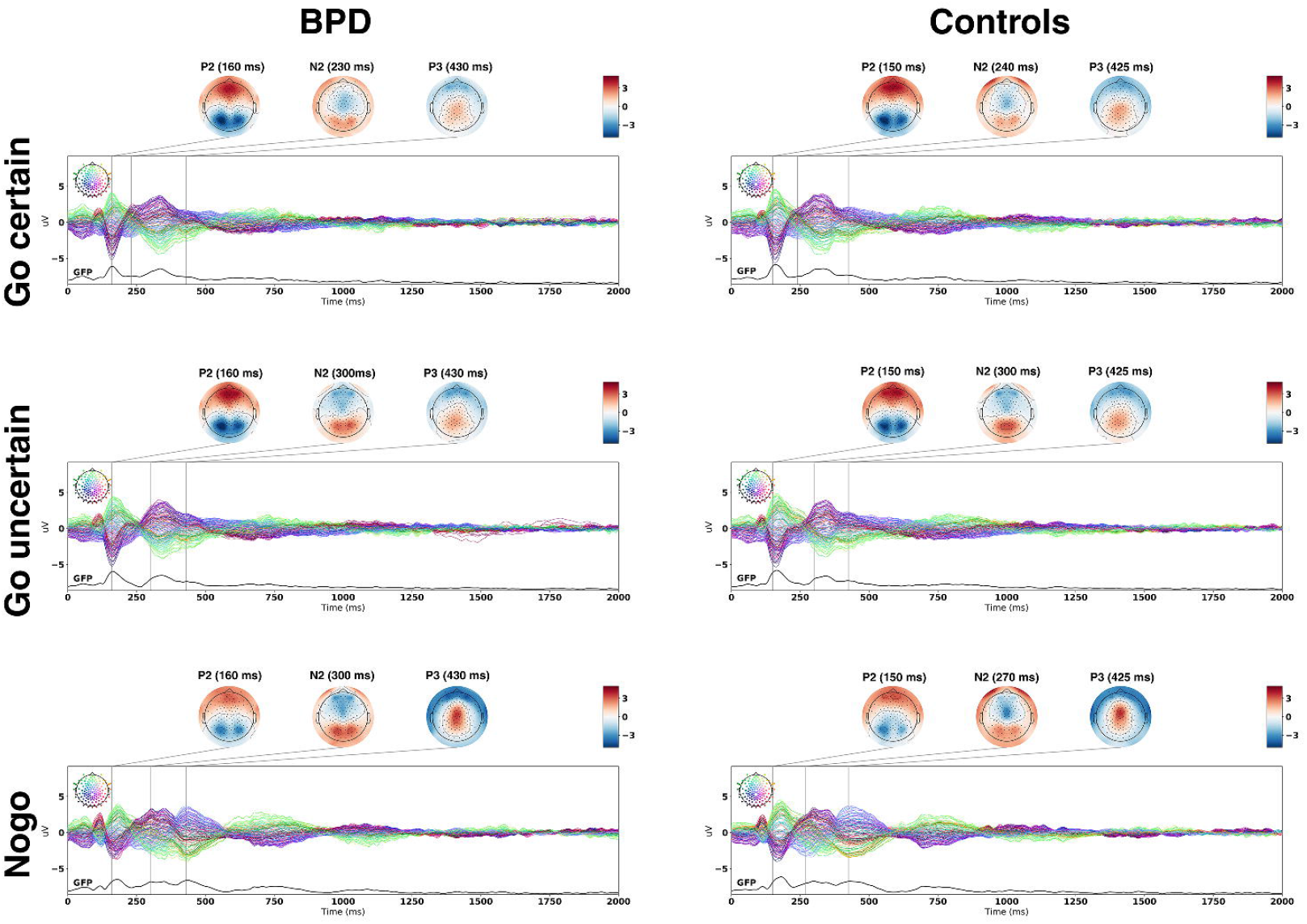

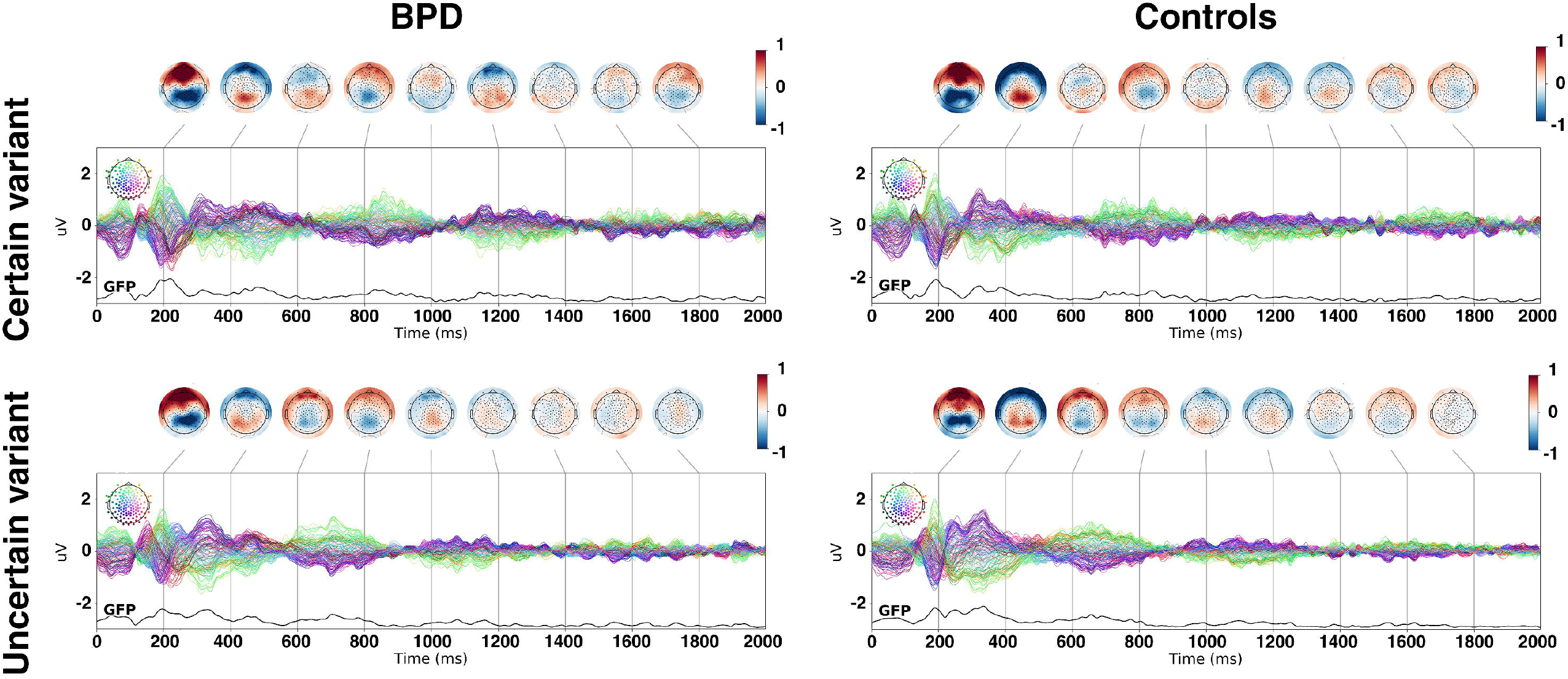

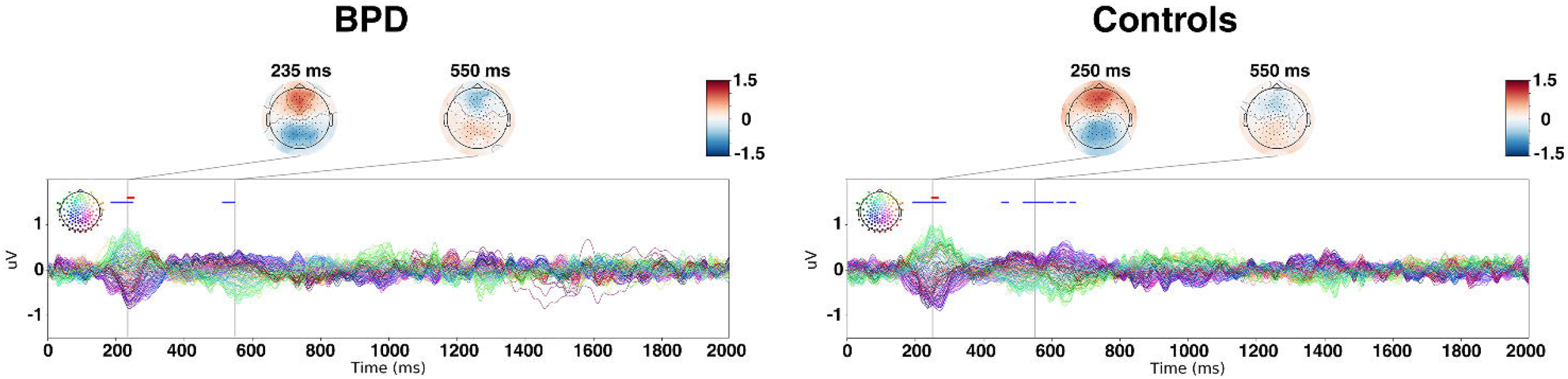

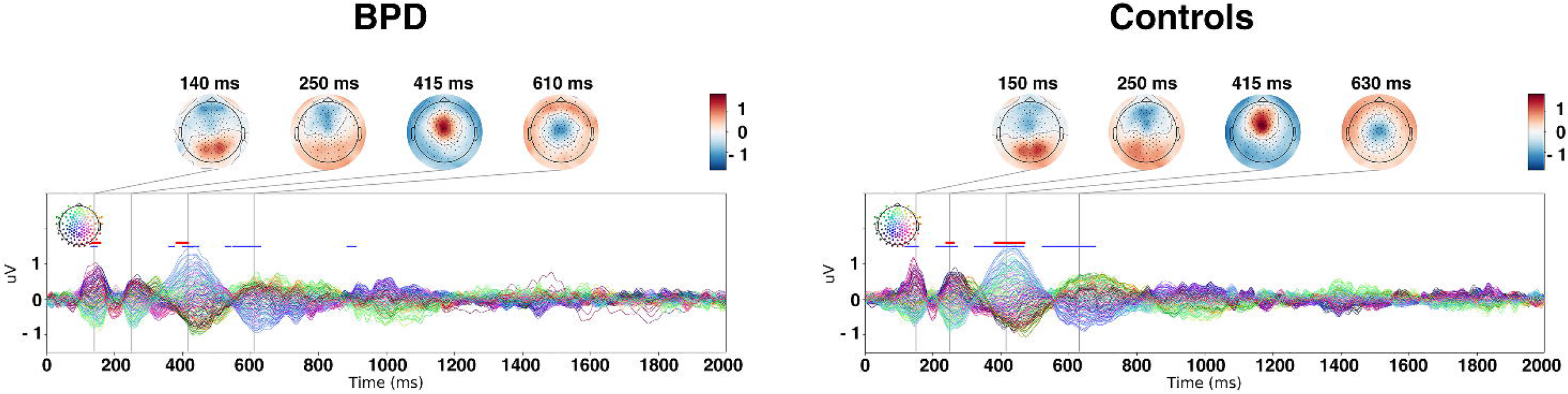

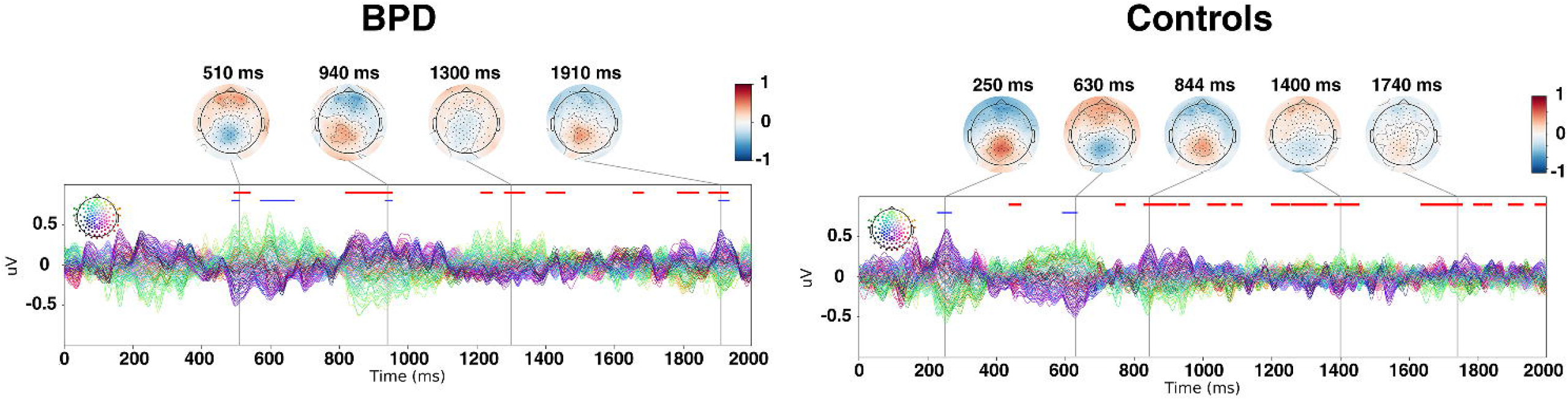

For the Go uncertain vs. Go certain condition comparison (Figure 7), both groups showed significant differences (p < 0.001), for both GFP and TANOVA measures, in the time-window of the N2 component (200 - 300 ms). In this time-window, both groups showed lower frontal negativity and lower occipital positivity in the Go uncertain condition. Besides, the N2 peaked later over the central areas in the Go uncertain compared to the Go certain condition (Figure 4). Furthermore, both groups showed significant differences in the TANOVA (p < 0.001), starting 453 ms after the stimulus onset for the healthy control group and 512 ms after the stimulus onset for the BPD group. No significant differences were observed in the time-windows of the P2 (140 - 250ms) and P3 (260 - 450ms) components.

For the NoGo condition vs. Go uncertain condition comparison (Figure 8), both groups showed significant differences (p < 0.001) in the time-windows of several components. In the time-window of the P2 component (140 - 250 ms), both groups showed lower frontal positivity and lower occipital negativity in the NoGo condition. Significant differences were observed in the GFP and TANOVA measures for the BPD group and only in the TANOVA for the HC group. In the time-window of the N2 component (200 - 300 ms), both groups showed the same tendency in amplitude differences, i.e. higher frontal negativity and higher occipital positivity in the NoGo condition. Significant differences (p < 0.001) in the GFP and TANOVA measures were observed in the HC group but not in the BPD group. In the time-window of the P3 component (260 - 450 ms), both groups showed higher fronto-central positivity in the NoGo condition. Significant differences (p < 0.001) in the GFP and TANOVA measures were observed in both groups. In the post-response period, we observed significant differences (p < 0.001) in the TANOVA both in the BPD (546 - 630 ms and 884 - 910 ms post-stimulus) and HC (522 - 677 ms post-stimulus) groups.

In both groups, lower activity in the late phase of the prestimulus period was observed in the uncertain compared to certain task variants (Figure 6). The permutation analysis on the GFP (Figure 9) revealed several intervals of significant differences within the 496 - 1930 ms and 438 - 2000 ms periods in the BPD and HC groups, respectively. In addition, the TANOVA revealed intervals of statistically significant differences within the 488 - 1933 ms and 228 - 634 ms periods in the BPD and HC groups, respectively.

In summary, we found a lower N2 with different topography in the Go uncertain compared to Go certain condition in both groups. We found a lower P2 in the NoGo condition compared to Go uncertain condition in the BPD group and the same trend in amplitude differences in the HC group. Additionally, in both groups, we observed topographical differences in the P2 component. We found a higher N2 with different topography in the NoGo condition compared to Go uncertain condition in the HC group. The same trend in N2 amplitude was observed in the BPD group. We found a higher P3 with different topography in the NoGo condition compared to Go uncertain condition in both groups. Finally, in both groups, lower activity in the uncertain compared to the certain task variant occurred predominantly later than 800 ms after the fixation cross presentation.

## 4. Discussion

Using high-density EEG we examined electrophysiological activity during a modified Go/NoGo task to search for correlates of proactive and reactive inhibitions in clinically impulsive BPD patients and in HC subjects. By contrasting the data within different task variants, conditions, and groups, the following key electrophysiological findings were revealed: 1) Lower GFP activity in the uncertain compared to certain task variant; 2) Lower N2 amplitude in the Go uncertain compared to Go certain condition; 3) Lower P2, higher N2, and higher P3 amplitudes in the NoGo condition compared to Go uncertain condition; 4) No electrophysiological difference between the BPD and HC groups. In addition to the electrophysiological findings, we observed longer reaction time in the Go uncertain compared to Go certain condition in both groups and lower accuracy in the Go uncertain condition in the BPD compared to HC group.

The following crucial mental processes were expected to occur successively during the employed cognitive task. During the prestimulus period in the certain task variant, the subject first decided to move, waited for the stimulus to occur, and then performed the motor action. In the prestimulus period of the uncertain task variant, however, the subject was both preparing for and proactively inhibiting motor action. In the early phase of the poststimulus period in the uncertain task variant the subject continued to inhibit the motor action, until the stimulus detection, stimulus discrimination, and decision “what to do” was finished. Consequently, in the late phase, the subject either performed or reactively inhibited the motor action. With these assumptions regarding the course of mental operations during the task our findings could be interpreted in terms of proactive and reactive inhibitions as outlined in the following section.

### 4.1 Proactive inhibition

In the later phase of the prestimulus period of both groups, we found lower GFP activity in the uncertain compared to certain task variants. Angelini et al. (2016) argues that the prestimulus activity is sensitive to cue-elicited motor priming that speeds up a subsequent motor response. The lower GFP activity observed here could therefore reflect reduced motor priming as a mechanism of proactive inhibition. Proactive inhibition was also suggested to be implemented via activation of the stopping network prior to the stimulus occurrence (Elchlepp et al., 2016). The authors argue that this activation is much weaker during proactive compared to reactive inhibition. The absence of an activity specific for the uncertain task variant in our study might be, therefore, explained by relatively weak involvement of the stopping network during the prestimulus period.

In the poststimulus period of both groups, we found initially almost identical evoked activity in the Go certain and Go uncertain conditions, indicating similar visual target processing in both conditions. Starting around 200ms after the stimulus onset, we found differences in the N2 component between the two conditions in both groups. Specifically, we found lower activity in the N2 time-window in the Go uncertain condition compared to Go certain condition together with differences in scalp topographies. In addition, we observed a later N2 peak over the central areas in the Go uncertain condition in both groups. The N2 was previously associated with both reactive inhibition (Kaiser et al., 2003; Bokura et al., 2001; Baumeister et al., 2014) and conflict monitoring (Donkers and van Boxtel, 2004; Nieuwenhuis et al., 2003) in the Go/NoGo context. A more recent fMRI-EEG study suggests N2 is associated with premotor activity originating from the supplementary and cingulate motor areas (Russo et al., 2016). Building on these findings, Perri (2020) suggested decreased N2 in successful Go trials compared to NoGo commissions to reflect suppressed premotor activity associated with proactive inhibition. In this frame, we suggest that the differences observed here in the N2 component might be caused by the differences in the timing of the decision making. In the Go certain condition the decision “what to do” took its course as soon as the fixation cross significance was identified and had already been finished prior to the stimulus onset. That is why the motor response can be initiated right after the stimulus detection. In the Go uncertain condition, the motor response still needs to catch up due to further processing costs imposed by the proactive inhibition that lasts until the decision “what to do” has been made. The decreased N2 in the Go uncertain condition might reflect suppressed premotor and motor processes as a mechanism of proactive inhibition. This view is supported by the longer reaction time as well as a later peaking N2 over the central areas in the Go uncertain condition.

To sum up, we propose the proactive inhibition in the modified Go/NoGo task is reflected in modulation of the late prestimulus activity and the poststimulus N2 activity. Our findings are in line with the view that proactive inhibition might be implemented via the activation of the stopping network prior to the stimulus onset and via reduced motor priming.

### 4.2 Reactive inhibition

In the poststimulus period of both groups, we observed differences in the P2, N2, and P3 components when comparing the NoGo to the Go uncertain condition. We found lower activity in the P2 component in the NoGo compared to the Go uncertain condition together with differences in scalp topographies. The P2 is an early wave occurring 150-220ms after the stimulus onset (Smith et al., 2004). Given its relatively short latency, we supposed that the P2 could reflect processes related to stimulus detection and discrimination rather than processes related to reactive inhibition. Benikos et al. (2013) found the P2 to be sensitive to task difficulty. Specifically, the authors found the P2 amplitude to be inversely related to the task difficulty. This relationship was much stronger for the NoGo than the Go condition, leading to significant differences between NoGo and Go P2 in the medium and hard task variants. Thus, the lower NoGo P2 in our study might reflect at least moderate task difficulty. The P2 was previously associated with sensory gating during the Go/NoGo task (Steele et al., 2014 ; Lijffijt et al., 2009) and was found to be correlated with alpha-inhibitory activity (Baijot et al., 2017). The alpha inhibition theory (Jensen and Mazaheri, 2010) proposes that the activity within the 8-13Hz range reduces the processing capacity in a given cortical area. It acts as a sort of “gating mechanism” for the optimal employment of sensory processing. Although the P2 is relatively rarely studied in the Go/NoGo context, this component seems to be highly relevant for visual attention and visual target processing. In line with other studies, we propose that the observed difference between Go uncertain and NoGo P2 amplitude might represent a visual gating mechanism, signifying increased processing demands in the NoGo condition.

We found higher N2 and P3 activities in the NoGo condition compared to the Go uncertain condition. The N2 component was previously associated both with early cognitive processes necessary for implementing inhibitory control (Luijten et al., 2014) and with the actual process of reactive inhibition (Kaiser et al., 2003). Alternatively, N2 is interpreted as a correlate of conflict monitoring (Donkers and van Boxtel, 2004). In a Go/NoGo task, a conflict might arise when the less frequent NoGo stimulus occurs because the more frequent stimulus is expected. Since in our study, the stimulus occurrences in the uncertain task variant were equiprobable, we suppose no such conflict arose. Therefore, the observed differences in the N2 component could rather account for premotor activity in the Go uncertain condition and/or for the processes related to reactive inhibition in the NoGo condition. Many previous studies have reported the P3 to be associated with reactive inhibition (Kenemans, 2015; Kok et al., 2004; Dimoska et al., 2006; Band and van Boxtel; 1999). Both of our groups had a strong P3 response to the NoGo stimuli. Our study thus supports the role of P3 in reactive inhibition.

To sum up, we propose the reactive inhibition in the modified Go/NoGo task is reflected in a modulation of the N2 and P3 components. In addition, our findings support the view that the P2 component is related to early visual processing rather than reactive inhibition processes.

### 4.3 Normal processing of emotionally neutral stimuli in borderline patients

Previous studies (Albert et al., 2019; Ruchsow et al., 2008) have found attenuated event-related activity in BPD patients compared to healthy subjects, particularly in the P3 component. Even though the BPD patients in the current study reported higher impulsivity on the UPPS-P scale, we observed no electrophysiological between-group differences related to proactive or reactive inhibition.

Recent studies suggest different processing of stressful stimuli in BPD compared to healthy subjects (Cackowski et al., 2014; Krause-Utz et al., 2016a). Hence, the electrophysiological activity in BPD patients might be modulated by stress. Our data provide evidence for the normal processing of emotionally neutral stimuli in BPD patients, suggesting that impulse control impairment in BPD might act through different mechanisms other than impaired proactive or reactive inhibition in neutral conditions. Similarly, we observed no relevant behavioral between-group differences. Both groups had similar response times and their accuracy exceeded 80%. Previous studies comparing BPD with healthy subjects found no differences (Völlm et al., 2004), as well as faster motor performance and higher error rates in BPD subjects (Hagenhoff et. al, 2013; Rentrop et al., 2008). We observed lower accuracy in BPD patients only in the Go uncertain condition but not in the Go certain or NoGo conditions. This might reflect problems related to attentional control, rather than proactive or reactive inhibition.

### 4.4 Limitations

The equal probability of the three conditions was used to eliminate the influence of the rarity of the stimuli. This might have led, however to a smaller electrophysiological response in the NoGo condition (Wessel, 2018) and therefore to less detectable correlates of reactive inhibition.

To find electrophysiological correlates of proactive and reactive inhibition we employed the previously used modification of the Go/NoGo task (Albares et al., 2014). This slow-paced and emotionally neutral task proved to be insensitive in detecting potentially existing electrophysiological differences between BPD and HC subjects. The Borderline profile, in general, is tied to problems in affective processing.

Studies suggest that impulsivity in borderline personality disorder is impacted by stress (Cackowski et al. 2014; Krause-Utz et al., 2016b). Therefore, a stressful or emotionally demanding Go/NoGo task might be required to reveal differences in proactive and reactive inhibition processing between BPD and a healthy population.

## 5. Conclusion

In conclusion, the present study describes the possible electrophysiological correlates of proactive and reactive inhibitions during a modified Go/NoGo task.

Our results suggest the proactive inhibition to be reflected in a modulation of the late prestimulus activity and suppression of the N2 component. Furthermore, we suggest that the reactive inhibition could be reflected in the enhancement of the N2 and P3 components. Our data provide evidence for the normal processing of emotionally neutral stimuli in BPD patients, suggesting that impulse control impairment in BPD might act through different mechanisms other than altered inhibitory control in an emotionally neutral environment.

## 6. Declarations

### Ethics approval and consent to participate

All participants gave their written informed consent prior to the experiment and the study received the approval of the Ethics Committee of University Hospital Brno. All experiments in this study were performed in accordance with the relevant guidelines and regulations.

### Consent for publication

Each author has read the complete manuscript and concurs with its content and publication.

### Availability of Data and Material

The data and material that support the findings of this study are available from the corresponding author upon reasonable request.

### Code availability

The custom in-house MATLAB code is available from the corresponding author upon reasonable request. RAGU software by Thomas Koenig is freely available http://www.thomaskoenig.ch/index.php/work/ragu.

### Author Contributions

J.K. pre-processed and analyzed the data, and wrote the initial draft of the manuscript. A.D. supervised the study, served as a consultant for clinical issues, wrote the initial draft of the manuscript. P.L. designed the study and collected the data. M.L. served as a consultant for the data analysis. B.J. and D.R. collected the data. M.B. and T.K. developed the initial idea. All authors reviewed the manuscript.

## Acknowledgements

The authors wish to thank Alexander Floyd for the proofreading assistance.

## References

Albares M, Lio G, Criaud M, Anton JL, Desmurget M, Boulinguez P (2014) The dorsal medial frontal cortex mediates automatic motor inhibition in uncertain contexts: Evidence from combined fMRI and EEG studies. Human Brain Mapping 35(11):5517–5531, DOI :10.1002/hbm.22567, URL https://dx.doi.org/10.1002/hbm.22567

Albert J, López-Martín S, Arza R, Palomares N, Hoyos S, Carretié L, Díaz-Marsá M, Carrasco JL (2019) Response inhibition in borderline personality disorder: Neural and behavioral correlates. Biological Psychology 143:32–40, DOI 10.1016/j.biopsycho.2019.02.003, URL https://dx.doi.org/10.1016/j.biopsycho.2019.02.003

Angelini M, Calbi M, Ferrari A, Sbriscia-Fioretti B, Franca M, Gallese V, Umiltà MA (2016) Proactive Control Strategies for Overt and Covert Go/NoGo Tasks: An Electrical Neuroimaging Study. PLOS ONE 11(3):e0152188–e0152188, DOI 10.1371/journal.pone.0152188, URL https://dx.doi.org/10.1371/journal.pone.0152188

Baijot S, Cevallos C, Zarka D, Leroy A, Slama H, Colin C, Deconinck N, Dan B, Cheron G (2017) EEG Dynamics of a Go/Nogo Task in Children with ADHD. Brain Sciences 7(12):167–167, DOI 10.3390/brainsci7120167, URL https://dx.doi.org/10.3390/brainsci7120167

Bailey NW, Hoy KE, Maller JJ, Segrave RA, Thomson R, Williams N, Daskalakis ZJ, Fitzgerald PB (2014) An exploratory analysis of go/nogo event-related potentials in major depression and depression following traumatic brain injury. Psychiatry Research: Neuroimaging 224(3):324–334, DOI 10.1016/j.pscychresns.2014.09.008, URL https://dx.doi.org/10.1016/j. pscychresns.2014.09.008

Band GPH, van Boxtel GJM (1999) Inhibitory motor control in stop paradigms: review and reinterpretation of neural mechanisms. Acta Psychologica 101(2-3):179–211, DOI 10.1016/s0001-6918(99)00005-0, URL https://dx.doi.org/10.1016/s0001-6918(99)00005-0

Barteček R, Hořínková J, Linhartová P, Kašpárek T (2019) Emotional impulsivity is connected to suicide attempts and health care utilization in patients with borderline personality disorder. DOI 10.1016/j.genhosppsych.2018.11.008, URL https://dx.doi.org/10.1016/j.genhosppsych.2018.11.008

Baumeister S, Hohmann S, Wolf I, Plichta MM, Rechtsteiner S, Zangl M, Ruf M, Holz N, Boecker R, Meyer-Lindenberg A, Holtmann M, Laucht M, Banaschewski T, Brandeis D (2014) Sequential inhibitory control processes assessed through simultaneous EEG–fMRI. NeuroImage 94:349–359, DOI 10.1016/j.neuroimage.2014.01.023, URL https://dx.doi.org/10.1016/j. neuroimage.2014.01.023

Benikos N, Johnstone SJ, Roodenrys SJ (2013) Varying task difficulty in the Go/Nogo task: The effects of inhibitory control, arousal, and perceived effort on ERP components. International Journal of Psychophysiology 87(3):262–272, DOI 10.1016/j.ijpsycho.2012.08.005, URL https://dx.doi.org/10.1016/j.ijpsycho.2012.08.005

Berchio C, Piguet C, Michel CM, Cordera P, Rihs TA, Dayer AG, Aubry JM (2017) Dysfunctional gaze processing in bipolar disorder. NeuroImage: Clinical 16:545–556, DOI 10.1016/j.nicl.2017.09.006, URL https://dx.doi.org/10.1016/j.nicl.2017.09.006

Black DW, Blum N, Pfohl B, Hale N (2004) Suicidal Behavior in Borderline Personality Disorder: Prevalence, Risk Factors, Prediction, and Prevention. Journal of Personality Disorders 18(3):226–239, DOI 10.1521/pedi.18.3.226.35445, URL https://dx.doi.org/10.1521/pedi.18.3.226.35445

Bokura H, Yamaguchi S, Kobayashi S (2001) Electrophysiological correlates for response inhibition in a Go/NoGo task. Clinical Neurophysiology 112(12):2224–2232, DOI 10.1016/s1388-2457(01)00691-5, URL https://dx.doi.org/10.1016/s1388-2457(01)00691-5

Braver TS (2012) The variable nature of cognitive control: a dual mechanisms framework. Trends in Cognitive Sciences 16(2):106–113, DOI 10.1016/j.tics.2011.12.010, URL https://dx.doi.org/10.1016/j.tics.2011.12.010

Brodsky BS, Malone KM, Ellis SP, Dulit RA, Mann JJ (1997) Characteristics of Borderline Personality Disorder Associated With Suicidal Behavior. American Journal of Psychiatry 154(12):1715–1719, DOI 10.1176/ajp.154.12.1715, URL https://dx.doi.org/10.1176/ajp.154.12.1715

Cackowski S, Reitz AC, Ende G, Kleindienst N, Bohus M, Schmahl C, Krause-Utz A (2014) Impact of stress on different components of impulsivity in borderline personality disorder. Psychological Medicine 44(15):3329–3340, DOI 10.1017/s0033291714000427, URL https://dx.doi.org/10.1017/s0033291714000427

Castro-Meneses, L. J., Johnson, B. W., & Sowman, P. F. (2015). The effects of impulsivity and proactive inhibition on reactive inhibition and the go process: Insights from vocal and manual stop signal tasks. Frontiers in Human Neuroscience, 9(OCT) doi:10.3389/fnhum.2015.00529

Cyders, M. A., Combs, J., Fried, R. E., Zapolski, T. C. B., & Smith, G. T. (2009). Emotion-based impulsivity and its importance for impulsive behavior outcomes. Impulsivity: Causes, control and disorders, 105–125.

Cyders MA, Smith GT (2007) Mood-based rash action and its components: Positive and negative urgency. Personality and Individual Differences 43(4):839–850, DOI 10.1016/j.paid.2007.02.008, URL https://dx.doi.org/10.1016/j.paid.2007.02.008

Delorme A, Makeig S (2004) EEGLAB: an opensource toolbox for analysis of single-trial EEG dynamics including independent component analysis. Journal of Neuroscience Methods 134(1):9–21, DOI 10.1016/j.jneumeth.2003.10.009, URL https://dx.doi.org/10.1016/j.jneumeth.2003.10.009

Dimoska A, Johnstone SJ, Barry RJ (2006) The auditory-evoked N2 and P3 components in the stop-signal task: Indices of inhibition, response-conflict or error-detection? Brain and Cognition 62(2):98–112, DOI 10.1016/j.bandc.2006.03.011, URL https://dx.doi.org/10.1016/j.bandc.2006.03.011

Donkers FC, van Boxtel GJ (2004) The N2 in go/no-go tasks reflects conflict monitoring not response inhibition. Brain and Cognition 56(2):165–176, DOI 10.1016/j.bandc.2004.04.005, URL https://dx.doi.org/10.1016/j. bandc.2004.04.005

Elchlepp H, Lavric A, Chambers CD, Verbruggen F (2016) Proactive in-hibitory control: A general biasing account. Cognitive Psychology 86:27–61, DOI 10.1016/j.cogpsych.2016.01.004, URL https://dx.doi.org/10.1016/j.cogpsych.2016.01.004

Galdo-Alvarez S, Bonilla FM, González-Villar AJ, de-la Peña MTC (2016) Functional Equivalence of Imagined vs. Real Performance of an Inhibitory Task: An EEG/ERP Study. Frontiers in Human Neuroscience 10:467–467, DOI 10.3389/fnhum.2016.00467, URL https://dx.doi.org/10.3389/fnhum.2016.00467

Grant BF, Chou SP, Goldstein RB, Huang B, Stinson FS, Saha TD, Ruan WJ (2008) Prevalence, correlates, disability, and comorbidity of DSM-IV borderline personality disorder: results from the Wave 2 National Epidemiologic Survey on Alcohol and Related Conditions. The Journal of clinical psychiatry 69(4):533–545

Habermann M, Weusmann D, Stein M, Koenig T (2018) A Student’s Guide to Randomization Statistics for Multichannel Event-Related Potentials Using Ragu. Frontiers in Neuroscience 12:355–355, DOI 10.3389/fnins.2018.00355, URL https://dx.doi.org/10.3389/fnins.2018.00355

Hagenhoff M, Franzen N, Koppe G, Baer N, Scheibel N, Sammer G, Gallhofer B, Lis S (2013) Executive functions in borderline personality disorder. Psychiatry Research 210(1):224–231, DOI 10.1016/j.psychres.2013.05.016, URL https://dx.doi.org/10.1016/j.psychres.2013.05.016

Jensen O, Mazaheri A (2010) Shaping Functional Architecture by Oscillatory Alpha Activity: Gating by Inhibition. Frontiers in Human Neuroscience 4:186–186, DOI 10.3389/fnhum.2010.00186, URL https://dx.doi.org/10.3389/fnhum.2010.00186

Kaiser S, Unger J, Kiefer M, Markela J, Mundt C, Weisbrod M (2003) Executive control deficit in depression: event-related potentials in a Go/Nogo task. Psychiatry Research: Neuroimaging 122(3):169–184, DOI 10.1016/s0925-4927(03)00004-0, URL https://dx.doi.org/10.1016/s0925-4927(03)00004-0

Kenemans JL (2015) Specific proactive and generic reactive inhibition.Neu-roscience & Biobehavioral Reviews 56:115–126, DOI 10.1016/j.neubiorev.2015.06.011, URL https://dx.doi.org/10.1016/j.neubiorev.2015.06.011

Koenig, T., Kottlow, M., Stein, M., & Melie-García, L. (2011). Ragu: a free tool for the analysis of EEG and MEG event-related scalp field data using global randomization statistics. Computational intelligence and neuroscience, 2011.Kok A, Ramautar JR, Ruiter MBD, Band GPH, Ridderinkhof KR (2004) ERP components associated with successful and unsuccessful stopping in a stop-signal task. Psychophysiology 41(1):9–20, DOI 10.1046/j.1469-8986.2003.00127.x, URL https://dx.doi.org/10.1046/j.1469-8986.2003.00127.x

Krause-Utz A, Cackowski S, Daffner S, Sobanski E, Plichta MM, Bohus M, Ende G, Schmahl C (2016a) Delay discounting and response disinhibition under acute experimental stress in women with borderline personality disorder and adult attention deficit hyperactivity disorder. Psychological Medicine 46(15):3137–3149, DOI 10.1017/s0033291716001677, URL https://dx.doi.org/10.1017/s0033291716001677

Krause-Utz A, Cackowski S, Daffner S, Sobanski E, Plichta MM, Bohus M, Ende G, Schmahl C (2016b) Delay discounting and response disinhibition under acute experimental stress in women with borderline personality disorder and adult attention deficit hyperactivity disorder. Psychological Medicine 46(15):3137–3149, DOI 10.1017/s0033291716001677, URL https://dx.doi.org/10.1017/s0033291716001677

Lehmann D, Skrandies W (1980) Reference-free identification of components of checkerboard-evoked multichannel potential fields. Electroencephalography and Clinical Neurophysiology 48(6):609–621, DOI 10.1016/0013-4694(80)90419-8, URL https://dx.doi.org/10.1016/0013-4694(80)90419-8

Lenzenweger MF, Lane MC, Loranger AW, Kessler RC (2007) DSM-IV Personality Disorders in the National Comorbidity Survey Replication. Biological Psychiatry 62(6):553–564, DOI 10.1016/j.biopsych.2006.09.019, URL https://dx.doi.org/10.1016/j.biopsych.2006.09.019

Liebrand M, Pein I, Tzvi E, Krämer UM (2017) Temporal Dynamics of Proactive and Reactive Motor Inhibition. Frontiers in Human Neuroscience 11:204–204, DOI 10.3389/fnhum.2017.00204, URL https://dx.doi.org/10.3389/fnhum.2017.00204

Lijffijt M, Lane SD, Meier SL, Boutros NN, Burroughs S, Steinberg JL, Moeller FG, Swann AC (2009) P50, N100, and P200 sensory gating: Relationships with behavioral inhibition, attention, and working memory. Psychophysiology 46(5):1059–1068, DOI 10.1111/j.1469-8986.2009.00845.x, URL https://dx.doi.org/10.1111/j.1469-8986.2009.00845.x

Linhartová P, širůček J, Ejova A, Barteček R, Theiner P, Kašpárek T (2021) Dimensions of Impulsivity in Healthy People, Patients with Borderline Personality Disorder, and Patients with Attention-Deficit/Hyperactivity Disorder. Journal of attention disorders 25(4):584–595

Links PS, Heslegrave R, van Reekum R (1999) Impulsivity: Core Aspect of Borderline Personality Disorder. Journal of Personality Disorders 13(1):1–9, DOI 10.1521/pedi.1999.13.1.1, URL https://dx.doi.org/10.1521/pedi.1999. 13.1.1

Luijten M, Machielsen M, Veltman D, Hester R, de Haan L, Franken I (2014) Systematic review of ERP and fMRI studies investigating inhibitory control and error processing in people. Journal of Psychiatry & Neuroscience 39(3):149–169, DOI 10.1503/jpn.130052, URL https://dx.doi.org/10.1503/jpn.130052

Michel CM, Murray MM (2012) Towards the utilization of EEG as a brain imaging tool. NeuroImage 61(2):371–385, DOI 10.1016/j.neuroimage.2011.12.039, URLhttps://dx.doi.org/10.1016/j.neuroimage.2011.12.039

Nguyen AT, Moyle JJ, Fox AM (2016) the N2 and the P3 modulation during partial inhibition in a modified GNG task. International Journal of Psychophysiology 107:63–71

Nieuwenhuis S, Yeung N, van den Wildenberg W, Ridderinkhof KR (2003) Electrophysiological correlates of anterior cingulate function in a go/no-go task: Effects of response conflict and trial type frequency. Cognitive, Affective, & Behavioral Neuroscience 3(1):17–26, DOI 10.3758/cabn.3.1.17, URL https://dx.doi.org/10.3758/cabn.3.1.17

Perri, R. L. (2020). Getting ready to act: Neurocognitive aspects of action preparation (Vol. 93). Sapienza Università Editrice.

Plesinger F, Jurco J, Halamek J, Jurak P (2016) SignalPlant: an open signal processing software platform. Physiological Measurement 37(7):N38–N48, DOI 10.1088/0967-3334/37/7/n38, URL https://dx.doi.org/10.1088/0967-3334/37/7/n38

Pompili M, Girardi P, Ruberto A, Tatarelli R (2005) Suicide in borderline personality disorder: A meta-analysis. DOI 10.1080/08039480500320025, URL https://dx.doi.org/10.1080/08039480500320025

Reeves-Dudley B (2017) Borderline Personality Disorder Psychological Treatment: An Integrative Review. Archives of Psychiatric Nursing 31(1):83–87, DOI 10.1016/j.apnu.2016.08.009, URL https://dx.doi.org/10.1016/j.apnu. 2016.08.009

Rentrop M, Backenstrass M, Jaentsch B, Kaiser S, Roth A, Unger J, Weisbrod M, Renneberg B (2008) Response Inhibition in Borderline Personality Disorder: Performance in a Go/Nogo Task. Psychopathology 41(1):50–57, DOI 10.1159/000110626, URL https://dx.doi.org/10.1159/000110626

Rihmer Z, Benazzi F (2010) Impact on suicidality of the borderline personality traits impulsivity and affective instability. Annals of clinical psychiatry: official journal of the American Academy of Clinical Psychiatrists 22:121–129

Roche RAP, Garavan H, Foxe JJ, O’Mara SM (2005) Individual differences discriminate event-related potentials but not performance during response inhibition. Experimental Brain Research 160(1):60–70, DOI 10.1007/s00221-004-1985-z, URL https://dx.doi.org/10.1007/s00221-004-1985-z

Ruchsow M, Groen G, Kiefer M, Buchheim A, Walter H, Martius P, Reiter M, Hermle L, Spitzer M, Ebert D, Falkenstein M (2008) Response inhibition in borderline personality disorder: event-related potentials in a Go/Nogo task. Journal of Neural Transmission 115(1):127–133, DOI 10.1007/s00702-007-0819-0, URL https://dx.doi.org/10.1007/s00702-007-0819-0

Russo FD, Lucci G, Sulpizio V, Berchicci M, Spinelli D, Pitzalis S, Galati G (2016) Spatiotemporal brain mapping during preparation, perception, and action. NeuroImage 126:1–14, DOI 10.1016/j.neuroimage.2015.11.036, URL https://dx.doi.org/10.1016/j.neuroimage.2015.11.036

Şahin M, Aybek E (2019) Jamovi: An Easy to Use Statistical Software for the Social Scientists. International Journal of Assessment Tools in Education 6(4):670–692, DOI 10.21449/ijate.661803, URL https://dx.doi.org/10.21449/ijate.661803

Sharif-Razi, M., Hodgins, D. C., & Goghari, V. M. (2019). Reactive and proactive control mechanisms of response inhibition in gambling disorder. Psychiatry Research, 272, 114-121. doi:10.1016/j.psychres.2018.12.049

Sheehan DV, Lecrubier Y, Sheehan KH, Amorim P, Janavs J, Weiller E, Dunbar, C G (1998) The Mini-International Neuropsychiatric Interview (MINI): the development and validation of a structured diagnostic psychiatric interview for DSM-IV and ICD-10. The Journal of clinical psychiatry

Skrandies W (1990) Global field power and topographic similarity. Brain Topography 3(1):137–141, DOI 10.1007/bf01128870, URL https://dx.doi.org/10.1007/bf01128870

Smith JL, Johnstone SJ, Barry RJ (2004) Inhibitory processing during the Go/NoGo task: an ERP analysis of children with attention-deficit/hyperactivity disorder. Clinical Neurophysiology 115(6):1320–1331, DOI 10.1016/j.clinph.2003.12.027, URL https://dx.doi.org/10.1016/j.clinph.2003.12.027

Soloff PH (2000) Characteristics of Suicide Attempts of Patients With Major Depressive Episode and Borderline Personality Disorder: A Comparative Study. American Journal of Psychiatry 157(4):601–608, DOI 10.1176/appi.ajp.157.4.601, URL https://dx.doi.org/10.1176/appi.ajp.157.4.601

Steele VR, Fink BC, Maurer JM, Arbabshirani MR, Wilber CH, Jaffe AJ, Sidz A, Pearlson GD, Calhoun VD, Clark VP, Kiehl KA (2014) Brain Potentials Measured During a Go/NoGo Task Predict Completion of Substance Abuse Treatment. Biological Psychiatry 76(1):75–83, DOI 10.1016/j.biopsych.2013.09.030, URL https://dx.doi.org/10.1016/j.biopsych.2013.09. 030

Sutton S, Baren M, Zubin J, John ER: Evoked-potential correlates of stimulus uncertainty. Science 150: 1187–1188, 1965.

Team RC (2020) R: A language and environment for statistical computing. R Foundation for Statistical Computing. Vienna, Austria, URL https://www.R-project.org/

Van Hulst, B. M., De Zeeuw, P., Vlaskamp, C., Rijks, Y., Zandbelt, B. B., & Durston, S. (2018). Children with ADHD symptoms show deficits in reactive but not proactive inhibition, irrespective of their formal diagnosis. Psychological Medicine, 48(15), 2508–2514. doi:10.1017/S0033291718000107

Völlm B, Richardson P, Stirling J, Elliott R, Dolan M, Chaudhry I, Ben CD, Mckier S, Anderson I, Deakin B (2004) Neurobiological substrates of antisocial and borderline personality disorder: preliminary results of a functional fMRI study. Criminal Behaviour and Mental Health 14(1):39–54, DOI 10.1002/cbm.559, URL https://dx.doi.org/10.1002/cbm.559

Wessel JR (2018) Prepotent motor activity and inhibitory control demands in different variants of the go/no-go paradigm. Psychophysiology 55(3):e12871–e12871, DOI 10.1111/psyp.12871, URL https://dx.doi.org/10.1111/psyp. 12871

Wessel JR, Aron AR (2015) It’s not too late: The onset of the frontocentral P3 indexes successful response inhibition in the stop-signal paradigm. Psychophysiology 52(4):472–480, DOI 10.1111/psyp.12374, URL https://dx.doi.org/10.1111/psyp.12374

Zandbelt BB (2011) Neural mechanisms of proactive and reactive inhibitory control: Studies in healthy volunteers and schizophrenia patients (Doctoral dissertation. Utrecht)

