## Supplementary material for "Electrophysiological correlates of proactive and reactive inhibition in a modified visual Go/NoGo task": Table 1

Table 1. The number of trials used for averaging (mean ± standard deviation) in different conditions separately for patient and control groups

| Characteristics | Borderline patients | Control subjects |
| --- | --- | --- |
|  | (n = 28) | (n = 35) |
| **Number of retained epochs**  Go uncertain  Go certain  No Go | 60.54 ± 5.69  53.04 ± 6.65  57.18 ± 7.48 | 61.71 ± 5.00  53.37 ± 7.55  59.00 ± 5.05 |
