## Supplementary material for "Electrophysiological correlates of proactive and reactive inhibition in a modified visual Go/NoGo task": Table 2

Table 2. Behavioral results

| Performance | BPD | HC | Test statistics | p-value |
| --- | --- | --- | --- | --- |
| parameters | (n = 28) | (n = 35) |  |  |
| **Mean reaction time ± SD (ms)** |  |  |  |  |
| Go uncertain condition | 375.1 ± 68.6 | 357.9 ± 46.5 | t (39.64) = 1.08 | 0.29 |
| Go certain condition | 317.3 ± 74.6 | 295.5 ± 44.5 | t (36.38) = 1.30 | 0.20 |
| **Mean accuracy ± SD (%)** |  |  |  |  |
| Go uncertain condition | 94.8 ± 5.6 | 98.5 ± 2.8 | W = 231.0 | **0.0019** |
| Go certain condition | 87.1 ± 9.4 | 88.9 ± 8.6 | W = 380.5 | 0.50 |
| NoGo condition | 88.4 ±7.5 | 91.0 ± 5.6 | W = 345.0 | 0.22 |

BPD = borderline personality disorder patients; HC = healthy controls; significant p-value (p<0.05) is highlighted in bold
