## Supplementary material for "Electrophysiological correlates of proactive and reactive inhibition in a modified visual Go/NoGo task": Figure caption

Figure 1.

Experimental protocol. Red fixation cross is followed by either Go or NoGo stimulus with equal probability. Green fixation cross is always followed by Go stimulus.

Figure 2.

Accuracy in three task conditions for both groups. Both groups have higher accuracy in Go uncertain condition compared to the other two conditions.

Figure 3.

Reaction times in Go certain and Go uncertain conditions for both groups. Both groups have longer response times in the Go uncertain compared to Go certain condition.

Figure 4.

Stimulus-locked grand average waveforms for the central cluster of electrodes, including FCz, for all three conditions. A) The evoked components (peak latency) in the borderline group: P2 (160 ms); N2 in the Go certain (230 ms), Go uncertain (300 ms), and NoGo (300 ms) conditions; P3 (430 ms). B) The evoked components (peak latency) in the healthy control group: P2 (150 ms); N2 in the Go certain (240 ms), NoGo (270 ms), and Go uncertain (300 ms) conditions; P3 (425 ms). Shadowed regions correspond to the 95% confidence interval bands. Scalp distributions of the evoked components at the peak latencies are presented in Figure 5.

Figure 5.

Stimulus-locked grand average waveforms in all electrodes for the Go certain, Go uncertain, and NoGo conditions in the borderline and healthy control groups. The topomaps show potential distributions at the latencies indicated in Figure 4.; GFP - Global field power.

Figure 6.

Fixation cross-locked grand average waveforms in all electrodes for the certain and uncertain task variants in the borderline and healthy control groups; GFP - Global field power. Note lower GFP with lower ERP amplitudes in the uncertain compared to certain task variant, e.g. at 1200ms, and 1800ms after the fixation cross onset, notably in the BPD group.

Figure 7.

Difference waveforms for Go uncertain minus Go certain conditions for both groups. The horizontal lines indicate intervals of statistically significant (p < 0.001) differences in the GFP (red) and TANOVA (blue) metrics. Significant differences were observed in both groups for both metrics within the time-window of the N2 component (note lower frontal negativity and lower occipital positivity in the Go uncertain condition at 235 ms and 250ms). Significant TANOVA differences were also observed in both groups in the late period (note topographical differences at 550 ms).

Figure 8.

Difference waveforms for NoGo minus Go uncertain conditions for both groups. The horizontal lines indicate intervals of statistically significant (p < 0.001) differences in the GFP (red) and TANOVA (blue) metrics. Significant differences were observed in both groups within the time-windows of the P2 (note lower frontal positivity and lower occipital negativity in the NoGo condition at 140 ms and 150 ms) and P3 (note higher fronto-central positivity in the NoGo condition at 415 ms) components, and in the late post-response period (note topographical differences at 610 ms and 630 ms). Significant differences were observed in the control group with the same tendency in the BPD group within the N2 component time-window (note higher frontal negativity and higher occipital positivity in the NoGo condition at 250 ms).

Figure 9.

Difference waveforms for uncertain minus certain task variants for both groups. The horizontal lines indicate intervals of statistically significant (p < 0.001) differences in the GFP (red) and TANOVA (blue) metrics; the topomaps illustrate topographical differences at several representative time points. The GFP permutation analysis revealed several intervals of significant differences within the 496 - 1930 ms and 438 – 2000 ms periods in the BPD and control groups, respectively. Besides, TANOVA revealed several intervals of significant differences within the 488 - 1933 ms and 228 - 634 ms periods in the BPD and control groups, respectively.
